# Multilayer network associations between the exposome and childhood brain development

**DOI:** 10.1101/2023.10.23.563611

**Authors:** Ivan L. Simpson-Kent, Mārtiņš M. Gataviņš, Ursula A. Tooley, Austin L. Boroshok, Cassidy L. McDermott, Anne T. Park, Lourdes Delgado Reyes, Joe Bathelt, M. Dylan Tisdall, Allyson P. Mackey

## Abstract

Growing up in a high poverty neighborhood is associated with elevated risk for academic challenges and health problems. Here, we take a data-driven approach to exploring how measures of children’s environments relate to the development of their brain structure and function in a community sample of children between the ages of 4 and 10 years. We constructed exposomes including measures of family socioeconomic status, children’s exposure to adversity, and geocoded measures of neighborhood socioeconomic status, crime, and environmental toxins. We connected the exposome to two structural measures (cortical thickness and surface area, *n* = 170) and two functional measures (participation coefficient and clustering coefficient, *n* = 130). We found dense connections *within* exposome and brain layers and sparse connections *between* exposome and brain layers. Lower family income was associated with thinner visual cortex, consistent with the theory that accelerated development is detectable in early-developing regions. Greater neighborhood incidence of high blood lead levels was associated with greater segregation of the default mode network, consistent with evidence that toxins are deposited into the brain along the midline. Our study demonstrates the utility of multilayer network analysis to bridge environmental and neural explanatory levels to better understand the complexity of child development.

## Introduction

Geography shapes socioeconomic opportunity (Massey et al., 2003). Neighborhood environments differ in income inequality, violence, and pollution, with consequences for children’s health and learning (Finkelhor et al., 2015; Manduca and Sampson, 2019; Spilsbury et al., 2006; Wodtke et al., 2022). Theories of how children’s experiences at home and in their neighborhoods influence their brain development range from specific to global (Rakesh and Whittle, 2021; Tooley et al., 2021, 2020). Specific theories focus on how individual brain regions are shaped by types of experiences: repeated experience of negative emotions could strengthen corticolimbic circuits that support emotion processing or regulation (Gee et al., 2013; Luby et al., 2017), and repeated experience of unpredictable threats could lead to vigilance and the strengthening of salience networks (McLaughlin et al., 2014). Global theories suggest that negative experiences and exposures cause physiological wear and tear (Brody et al., 2014; McEwen, 1998; Raffington et al., 2023), leading to broad changes in brain development, including accelerated structural development (Belsky, 2019; Tooley et al., 2021). Less is known about whether adversity is associated with accelerated functional development. Many studies have shown a less developmental change in functional brain measures in youth who have been exposed to adversity (Chahal et al., 2022; Park et al., 2021; Tooley et al., 2020), rather than earlier development. Specific and global theories have shaped experiments and analyses in the field, but they are not mutually exclusive, nor do they include all environmental influences. For example, psychosocial stressors often co-occur with environmental toxins like lead and air pollution, which also have been linked to structural and functional brain differences (Bahrami et al., 2022; Marshall et al., 2020; Reuben et al., 2020).

The majority of environmental neuroscience studies have used univariate approaches, studying individual exposures at a time (Liu et al., 2022). As a result the associations among multiple exposures, as well as between different exposures and the brain, cannot be identified. An alternative approach is to consider multiple exposures and to simultaneously model their relationships as an exposome (Wild, 2005). The exposome, defined as the totality of environmental exposures across the lifespan, was originally established to address the need for more comprehensive evaluation of environmental exposures in relation to epidemiological outcomes (Wild, 2012). Since then, psychiatry studies have investigated the exposome’s role in relation to psychopathological outcomes using factor analysis, which reduces multiple exposures to a few factors (e.g., Barzilay et al., 2021; Guloksuz et al., 2018; Moore et al., 2022; Pries et al., 2022; Sydnor et al., 2023). However, this approach alone might not be able to fully capture the complexity of relationships between environmental exposures and outcomes of interest. A complementary method is to model the exposome as a network, whereby specific environmental exposures can be conceptualized to interact with each other and influence not only mental health outcomes but also their biological underpinnings such as brain structure and function. This network approach for the exposome originates from a complex systems framework that conceptualizes psychopathology as arising from the interaction of individual symptoms (Borsboom, 2017; Robinaugh et al., 2020).

Once the exposome is constructed, it can be connected to measures of brain development using a multilayer network approach (Bianconi, 2018). The multilayer network approach provides a common framework to conceptualize and estimate the associations within the exposome, within the brain, and their connections between each other. One common use of multilayer network analyses is to model the conditional dependencies among variables across layers such as clinical outcomes and brain networks (Blanken et al., 2021). These statistical dependencies are often estimated using partial correlations, whereby any connection between two nodes represents a statistical relationship that remains after controlling for the associations between those two nodes with all other nodes in the network. This method has been used previously to characterize relationships among brain measures and cognitive and mental health outcomes (Bathelt et al., 2022; Hilland et al., 2020; Simpson-Kent et al., 2021).

In this study, we asked how children’s experiences in their homes and in their neighborhoods related to their brain structure and function. We recruited families from diverse areas of the Philadelphia, Pennsylvania region who varied along multiple dimensions including exposure to neighborhood violence, lead and pollution, and income inequality. Children between the ages of 4 and 10 years participated in structural and resting-state functional MRI, and their parents reported on family education, income, and adverse childhood experiences (ACEs). We geocoded neighborhood-level factors from family addresses. We constructed an exposome to relate children’s experiences and exposures and connected it to structural and functional brain phenotypes. For brain structure, we focused on cortical thickness and cortical surface area. Cortical thickness declines during development and cortical surface area first expands and then contracts (Bethlehem et al., 2022; Rutherford et al., 2022). For brain function, we used resting-state functional connectivity to model the cortex as a network (Bassett et al., 2018), and focused on two metrics of cortical network architecture that have been shown to change with development: the clustering coefficient, a measure of network segregation, and the participation coefficient, a measure of network integration (Guimerà and Amaral, 2005; Rubinov and Sporns, 2010). During development, cortical network architecture is refined, with increases in the clustering coefficient reflecting denser local connectivity between nearby nodes, and decreases in the participation coefficient reflecting fewer diverse connections between distant nodes (Cohen and D’Esposito, 2016; Fair et al., 2009, 2007; Marek et al., 2015; Tooley et al., 2022a, 2022b, 2020) To allow for comparability across measures, and to avoid overloading the models, we used a seven-system division of cortex into systems based on functional connectivity (Yeo et al., 2011). Our data-driven exploratory approach allows us to both interrogate existing theories of how children’s experiences shape their brains and to generate new insights into previously undescribed relationships.

## Methods

### Participants

This study was approved by the Institutional Review Board (IRB) at the University of Pennsylvania. Parents provided informed consent. Children below 8 years old provided verbal consent and children 8 years old and older provided written assent.

Children between the ages of 4 and 10 were recruited from Philadelphia, Pennsylvania and its surrounding areas through advertisements on public transportation and social media, outreach programs, partnerships with local schools, and community family events. Participants were screened prior to participation and were excluded if they had a diagnosis of a psychiatric, neurological, or learning disorder, were born more than six weeks premature, were adopted, or had any contraindications for MRI scanning. A total of 212 participants participated in scanning at their first session. When including all timepoints, 217 participants participated in scanning. Of these 217 participants, 211 scanned at their first session, 43 at their second session 4-42 months later (mean: 13.6, median: 12 months), and 8 at their third session (9-21 months after second).For structural analyses, participants were excluded if they had low-quality images or poor quality surface reconstructions (n = 42; see ***Structural MRI preprocessing***). 170 participants were included in the structural analyses (only data from the first time point were included). For functional analyses, which rely on resting-state data that are more difficult to collect in young children, we considered all possible time points and included the time point with the lowest framewise displacement (n = 217 participants, 262 scan sessions). Participants were excluded if they did not complete a resting state scan (n = 32 participants, 34 scan sessions), fell asleep during the scan (n = 11 participants, 14 scan sessions), had poor quality structural data for registration during preprocessing (n = 8 participants, 8 scan sessions), or had high in-scanner head motion (n = 26 participants, 31 sessions) or insufficient scan data from resting-state functional images (n = 10 participants, 12 sessions; see ***Resting-state functional MRI preprocessing*** for criteria). From the 163 sessions available for the 130 participants remaining, we picked the earliest available session, resulting in the final functional imaging dataset included 120 scans from the first time point, 9 at the second time point, and 1 at the third time point. Statistical comparison of the included and excluded subjects for both samples is available in **Supplemental Tables 1 and 2** for the structural and functional samples, respectively.

Parents were asked to report their child’s date of birth, gender, race, and ethnicity. Parents were provided with four options for gender: female, male, other, or prefer not to answer. We acknowledge that these response options do not fully reflect the array of gender identities and conflate sex and gender. Parents provided their child’s race out of the following options: Black, White, Asian, Native Hawaiian or Other Pacific Islander, American Indian or Alaska Native, and Other. They could indicate more than one race category. Additionally, for ethnicity, parents were asked if their child was Hispanic or Latinx. Parents reported their total annual family income in one of 11 income bins (less than $5,000; $5000-$11,999; $12,000-$15,999; $16,000-$24,999; $25,000-$34,999; $35,000-$49,999; $50,000-$74,999; $75,000-$99,999; $100,00-$149,999; $150,000-$199,999; and $200,000 or greater). For analytical purposes, income was re-coded to represent the median of each bin; the maximum possible income in our sample was $200,000. Demographic information, including these data, is provided in **Table 1**.

**Table 1.** Demographic and geocoded information of participants in both neuroimaging samples. *Survey form allowed parents to endorse more than one racial identity hence the sum of racial identities is greater than 100%. For comparison, Philadelphia’s population is 43.6% Black, 44.8% White, 7.8% Asian, 3.9% Other, and 15.2% Hispanic or Latino (US Census Bureau, 2020). ^十^Market Profile variables indices represent deviations from the U.S. average rate of that specific crime - U.S. mean rate of a specific crime is 100.

| | Structural imaging<br>($n = 170$) | Functional imaging<br>($n = 130$) |
| --- | --- | --- |
| Age (years, mean [SD] {range}) | 6.34 [1.40] {4.05-10.59} | 6.64 [1.35] {4.11-10.59} |
| Sex (n, Female/Male/Other) | 89 / 81 / 0 | 69 / 61 / 0 |
| Race* (%) | 50.6 % Black | 51.5 % Black |
|  | 41.8 % White<br>11.8 % Asian<br>1.8% Native Hawaiian or<br>Other Pacific Islander<br>1.8% American Indian or<br>Alaska Native<br>6.5% Other | 42.3 % White<br>13.8 % Asian<br>2.3% Native Hawaiian or<br>Other Pacific Islander<br>1.5% American Indian or<br>Alaska Native<br>3.1% Other |
| Ethnicity (%) | 11.3% Latinx/Hispanic | 10.9% Latinx/Hispanic |
| Family Income (thousands of \$, mean [SD]) | 80.1 [68.2] | 85.7 [68.2] |
| Parent Education (years, mean [SD]) | 14.88 [2.72] | 15.09 [2.72] |
| Adverse Childhood Experiences (ACEs)<br>(mean, [SD]) | 0.97 [1.35] | 1.02 [1.31] |
| Neighborhood unemployment, (% mean [SD]) | 34.7 [24.5] | 34.3 [23.9] |
| Percentage of people (25+ years old) with a<br>Bachelor's degree in the neighborhood (%<br>mean [SD]) | 8.3 [5.4] | 8.3 [5.4] |
| Percentage incidence of high blood levels in<br>neighborhood (% mean [SD]) | 6.3 [3.4] | 6.07 [3.4] |
| Average particulate matter concentration in<br>neighborhood ($\mu\text{g}/\text{m}^3$, mean [SD]) | 1.30 [0.33] | 1.32 [0.33] |
| Murder Index<sup>+</sup> (unitless, mean [SD]) | 110.5 [50.8] | 108.3 [49.2] |
| Rape Index<sup>+</sup> (unitless, mean [SD]) | 94.0 [53.0] | 91.5 [52.2] |
| Robbery Index<sup>+</sup> (unitless, mean [SD]) | 90.1 [58.2] | 88.4 [58.0] |
| Assault Index<sup>+</sup> (unitless, mean [SD]) | 113.2 [41.8] | 109.3 [45.0] |
| Larceny Index<sup>+</sup> (unitless, mean [SD]) | 115.2 [41.8] | 111.7 [42.2] |
| Burglary Index<sup>+</sup> (unitless, mean [SD]) | 112.1 [50.4] | 108.4 [52.2] |
| Gini Index (unitless, mean [SD]) | 0.460 [0.065] | 0.458 [0.063] |

Parents completed the 10-item Adverse Childhood Experiences (ACEs) questionnaire about their children. The ACEs questionnaire is a widely-used assessment of early childhood experiences, which measures sexual, physical and emotional abuse, witnessing domestic violence, physical and emotional neglect, parental separation or divorce, and substance abuse, mental illness, or incarceration of an adult in the household (Murphy et al., 2016). An ACEs score is calculated by summing the responses to each of the adversity categories, with a maximum possible score of 10. See **Table 1** for descriptive statistics of the ACEs questionnaire for our structural and functional neuroimaging samples.

### Geocoding

Participant addresses were geocoded on a secure network using an offline ArcGIS address locator dataset of addresses and coordinates across the US provided by the University of Pennsylvania Libraries using ArcGIS v10.7 (Esri Inc., 2019). The addresses were transformed to geospatial coordinates in Pennsylvania and New Jersey. One participant with an address in Virginia was excluded from geocoding analyses. Address values that could not be extracted either due to an incorrect street, house number, or post box entries were excluded from further geocoding steps (*n* = 21). Address coordinate values were mapped onto Census Tract (CT) Shapefile fields, taken from the U.S. Census Bureau Census Tract Map for 2019 (U.S. Census Bureau, 2019a), and then the census tract IDs were paired with the respective subject IDs. All geocoded demographic and environmental variables were extracted at the census tract level by pairing with the census tract corresponding to the living address of a participant. Geocoded measures capture information related to neighborhood crime levels, neighborhood socioeconomic status (SES), and pollution levels. For descriptions of all geocoded measures, see **Supplementary Table 3**. See **Table 1** for descriptive statistics of geocoded variables for our structural and functional neuroimaging samples.

Market Profile Data is a census-level aggregate of demographic data, such as crime, population, employment, and access to businesses and resources (Easy Analytic Software, Inc., 2020). We used the Market Profile crime indices which are predicted using a regression model previously trained to predict county-level crime data (Federal Bureau of Investigation crime data) from county-level demographics. EASI shows that these models are highly accurate at predicting county-level crime data based on demographic data, in a separate, test set of counties. These models trained on county-level data are, then, used to predict crime indices at the tract level using tract-level demographic data. The crime variables are quantified as deviations from the U.S. average where the U.S. average rate of a specific kind of crime is represented as 100 and raw deviations in crime rate are weighted by the population in the census tract. Violent crime categories across these measures are murder, aggravated assault, rape, and robbery, and non-violent crime categories include larceny and burglary, according to the Uniform Crime Report of the Federal Bureau of Investigation (Federal Bureau of Investigation, 2019). We used structural equation modeling to determine whether crime variables were best included in the network as a single, sum score node or as separate crime index nodes (see **Supplemental Methods** for analysis details).

Three neighborhood SES variables were drawn from the American Community Survey (ACS). We used the ACS 2015-2019 5-year release due to the increased accuracy of the results when the measures are sampled across a bigger number of entries per census tract (U.S. Census Bureau, 2019b). The unemployment measure was calculated by dividing the number of people unemployed in the labor force divided by the number of people in the labor force above the age of 16. The education measure was the number of people over 25 with an advanced degree (Bachelor’s degree or higher) divided by the number of people over the age of 25 in the census tract. The Gini Index was used to estimate income distribution inequality within the census tract. The Index is calculated by taking the difference between the actual distribution of income and the ideal distribution. An index of 0 indicates perfect equality (each person owns an equal number of resources) and 1 denotes perfect inequality (few people own the majority of the resources) (“Gini Index,” 2008). In 2021, the United States had a Gini index of 0.494, which can be compared to the households in the 90th income percentile earning 13.53 times more than those in the 10th percentile (Semega et al., 2022).

Particulate matter 2.5 (PM2.5) tract-level data were predicted from regression models trained to predict Environmental Protection Agency (EPA) sensors based on local environmental measures, such as land use and traffic (Kim et al., 2020; Wang et al., 2020). EPA sensor results were predicted using area-specific environmental measures and then these trained models were used to predict tract-level concentration data (Kim et al., 2020; Wang et al., 2020).

Child blood lead level is based on the Philadelphia Department of Public Health lead surveillance campaign “Lead and Healthy Homes program” (Lead and Safe Homes Program, 2019). Children whose blood lead levels are above 5 µg/mL are classified as having high blood lead levels. This number is scaled by the number of children tested in a specific census tract. The data used in this study comes from the 2013-2015 data sample as this is publicly available on OpenData Philly (City of Philadelphia Department of Public Health, 2017). This measure was only available for children living in Philadelphia County (*n* = 95 for structural analyses, *n* = 76 for functional analyses).

### Neuroimaging data acquisition

Imaging was performed at the Center for Advanced Magnetic Resonance Imaging and Spectroscopy (CAMRIS) at the University of Pennsylvania. Scans were conducted using a 3 Tesla MRI scanner (MAGNETOM Prisma, Siemens Healthineers, Erlangen, Germany) with the vendor’s 32-channel head coil. A whole-brain, high-resolution, T1-weighted 3D-encoded multi-echo structural scan (MEMPRAGE) was collected (acquisition parameters: TR = 2530 ms, TI = 1330 ms, TEs = 1.69 ms/3.55 ms/5.41 ms/7.27 ms, bandwidth (BW) = 650 Hz/px, 3x GRAPPA, flip angle = 7°, voxel size = 1 mm isotropic, matrix size = 256 × 256 ×176, FOV = 256 mm, total scan time = 4:38). This anatomic sequence used interleaved volumetric navigators to prospectively track and correct for subject head motion (Tisdall et al., 2012). One or two five-minute resting-state fMRI scans were acquired using a T2*-weighted multiband gradient-echo echoplanar imaging (EPI) sequence (acquisition parameters: TR = 2000 ms, TE = 30.2 ms, BW = 1860 Hz/pixel, flip angle = 90°, voxel size = 2 mm isotropic, matrix size = 96 × 96, 75 axial slices, FOV = 192 mm, volumes = 150–240, 5 dummy scans, multiband acceleration factor = 3). We chose a multiband factor of three to minimize interactions between multiband and motion (Risk et al., 2021).

### Structural MRI preprocessing

Structural image quality was manually assessed by two lab members individually, without knowledge of participant information and demographics. Ratings ranged from 1 (good quality) to 4 (poor quality), and were averaged across raters. Cortical surfaces were reconstructed using FreeSurfer 6.0.0 (Dale et al., 1999; Fischl, 2012). After surface reconstruction, structural images were inspected and edited for cortical surface reconstruction errors. In images rated above 3.5, if reconstruction errors could not be resolved, participants were excluded (*n* = 17). Morphometric measures (surface area and cortical thickness) were extracted from a 400-region parcellation (Schaefer et al., 2018). Parcel-wise cortical thickness was averaged within systems and surface area was summed within systems using the 7-system partition: visual, somatomotor, limbic, dorsal attention, ventral attention, executive control, and default mode systems (Yeo et al., 2011). “System” is used to refer to a set of regions defined *a priori* (e.g., dorsal attention, frontoparietal or control, default mode networks).

### Resting-state functional MRI preprocessing

Participant scans were excluded if average framewise displacement (FD) was greater than 1 mm or if more than 30% of the volumes had FD > 0.5 mm, and if the scan had fewer than 130 volumes. Data that passed motion criteria also passed visual inspection for whole-brain field of view coverage, signal blurring or artifacts, and proper alignment to the anatomic image. fMRIPrep visual reports, MRIQC version 0.14.2, and xcpEngine scan quality outputs were used for visual inspection. Preprocessing was performed using fMRIPrep version 1.2.6-1 (RRID:SCR_016216; Esteban, Markiewicz, et al., 2019; Esteban, Wright, et al., 2019), which is based on Nipype version 1.1.7 (RRID:SCR_002502; Gorgolewski et al., 2011; nipy/nipype:1.1.7); as well as xcpEngine version 1.0 (Ciric et al., 2018). Cortical surfaces were reconstructed using *recon-all* (Dale et al., 1999). T1-weighted (T1w) images were corrected for intensity nonuniformity using N4BiasFieldCorrection (Tustison et al., 2010; Advanced Normalization ToolS (ANTS) version 2.2.0), and used as T1w references throughout the workflow. The T1w image was skull stripped using *antsBrainExtraction.sh* script (ANTS version 2.2.0) using OASIS as the target template. The brain mask was refined with a custom variation of the method to reconcile ANTS-derived and FreeSurfer-derived segmentations of the cortical gray matter of Mindboggle (RRID:SCR_002438, Klein et al., 2017). Spatial normalization of the T1w image to the ICBM 152 Nonlinear atlases version 2009c (RRID:SCR_008796; Fonov et al., 2011) was performed through nonlinear registration with antsRegistration (ANTS version 2.2.0, RRID:SCR_004757; Avants et al., 2010). Brain tissue segmentation of CSF, WM, and gray matter was performed on the brain-extracted T1w using FAST [Functional MRI of the Brain Software Library (FSL) version 5.0.9; RRID:SCR_002823; Zhang et al., 2001]. For each of the resting-state BOLD runs, the following preprocessing steps were performed. A reference volume and its skull-stripped version were generated using a custom methodology of fMRIPrep. The BOLD reference was then coregistered to the T1w reference using *bbregister* (FreeSurfer) which implements boundary-based registration (Greve and Fischl, 2009). Coregistration was configured with nine degrees of freedom to account for distortions remaining in the BOLD reference. Head-motion parameters were estimated before spatiotemporal filtering using the mcflirt tool (FSL version 5.0.9; Jenkinson et al., 2002). Slice-time correction was done with 3dTshift from Analysis of Functional NeuroImages (AFNI) 20160207 (RRID:SCR_005927; Cox and Hyde, 1997). The functional images were resampled onto MNI152NLin2009cAsym standard space by applying a single, composite transform, generating a preprocessed BOLD run in MNI152NLin2009cAsym space.

We used a confound regression procedure that has been optimized to reduce the influence of participant motion, implemented in xcpEngine version 1.0 (Ciric et al., 2017; Parkes et al., 2018; Satterthwaite et al., 2013b), a multimodal tool kit that deploys processing instruments from frequently used software libraries, including FSL (Jenkinson et al., 2012) and AFNI (Cox, 1996). Functional time series were demeaned, linear and quadratic terms were removed, and confound regression was performed. Confound regression was done using a 36-parameter model which accounted for mean whole-brain signal, signal from WM and CSF compartments, six motion parameters, and their derivatives, quadratic terms, and quadratic terms of the derivatives (Satterthwaite et al., 2013a). Motion censoring was performed with the following criteria for removing time points: framewise displacement (FD) greater than 0.5 mm or standardized DVARS (root-mean-square intensity difference from one volume to the next) greater than 1.75. Outlier volumes were interpolated over using least-squares spectral analysis (Power et al., 2014) before bandpass filtering to retain frequencies between 0.01 Hz and 0.08 Hz, then censored again. Before confound regression, all confound parameters were bandpass filtered the same way as the original time series data, ensuring comparability of the signals in frequency content (Hallquist et al., 2013).

### Functional network analysis

As part of the xcpEngine pipeline, we extracted BOLD time series from preprocessed and nuisance-regressed data using the Schaefer400 cortical parcellation (Schaefer et al., 2018), which subdivides the Yeo 7 parcellation into evenly sized parcels. Functional connectivity matrices were averaged across scan runs during one scan session, weighted by the number of frames in each run passing the inclusion criteria.

Correlation matrices were represented as graphs or networks (Bassett et al., 2018), where regions are represented as network nodes and correlations between time series of pairs of regions are represented as the edges between nodes. Product-moment correlations were calculated between parcels *i* and *j* and represented the edge weights of the edges between region *i* and *j* (Zalesky et al., 2012). Fisher Z-transformation was performed on the functional connectivity network matrix (of edge weights). The final networks were unthresholded, signed, and weighted. We chose to use weighted rather than binary edges to reflect variation in the strength of connectivity (Cole et al., 2012; Rubinov and Sporns, 2011; Santarnecchi et al., 2014), and included both positive and negative edges because of evidence that anticorrelations are meaningful (Chai et al., 2014; Nenning et al., 2023; Santarnecchi et al., 2014).

We calculated two graph measures, participation and clustering coefficient, to characterize network integration and segregation. Participation and clustering coefficients were calculated at the nodal level and then averaged across the same 7-system partition as the structural MRI data: visual, somatomotor, limbic, dorsal attention, ventral attention, executive control, and default mode systems (Yeo et al., 2011). These. All calculations of graph measures were performed using MATLAB R2020b (The MathWorks Inc., 2020) and the Brain Connectivity Toolbox (Rubinov and Sporns, 2010).

The participation coefficient quantifies the extent of integration of a region across different systems. A higher participation coefficient reflects more diverse connections to different systems (Guimerà and Amaral, 2005; Rubinov and Sporns, 2010). The participation coefficient for a node *i* (*P_i_*) was calculated as follows:

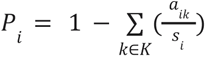

Where *k* is a system (e.g., dorsal attention) as part of the set *K* of all 7 systems (Yeo et al., 2011), *a_ik_* is the sum of all edge weights between node *i* and all of the nodes in *k*, and *s_i_* is the sum of all edge weights from node *i* (otherwise referred to as node strength). In these analyses of a signed network, the participation coefficient was calculated for positive and negative edges separately (as implemented by Rubinov and Sporns, 2010), then negative and positive-edge participation coefficients were averaged at the nodal level and then at the system level.

The clustering coefficient is a measure of local segregation that quantifies a node’s local neighborhood capacity to support information transfer (Achard et al., 2006; Bartolomei et al., 2006; Bassett et al., 2006; Tooley et al., 2022b, 2020; Xu et al., 2016). The clustering coefficient captures the strength of triangles of edges and nodes that surround a node. Nodes with high clustering coefficients show strong connections to nodes that also have strong connections to each other. We used an edge weight sign-sensitive generalization of the clustering coefficient calculation (Costantini and Perugini, 2014; Zhang and Horvath, 2005). It distinguishes between triangle signs (positive and negative triangles) and adjusts for nonredundancy in path information based on edge weight signs. For a node *i*, clustering coefficient, *C_i_*, is calculated as follows:

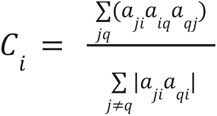

Where *i*, *j*, and *q* are distinct (neighboring) nodes with edge weights *a_ij_*between nodes *i* and *j*.

### Multilayer network creation

Statistical analyses were performed using R (R Core Team, 2023) version 4.2.3 (“Funny-Looking Kid”) and RStudio version 2022.07.2 (“Spotted Wakerobin”). Summary statistics and plots of networks were conducted using the packages *corrplot* (version 0.92, Wei and Simko, 2021), *jmv* (version 2.3.4, Selker et al., 2022), *OpenMx* (version 2.21.1, Boker et al., 2023; Neale et al., 2016), *psych* (version 2.2.5, Revelle, 2022), *reshape2* (version 1.4.4, Wickham, 2007), *gtsummary* (version 1.7.2, Sjoberg et al., 2021), and *dplyr*, *tidyr*, and *ggplot2* from tidyverse (Wickham et al., 2019).

Because age and motion are significantly related to many of the brain measures studied here, we residualized all brain measures for linear effects of age and scan quality measures. Structural measures were residualized for linear effects of age and manually-rated scan quality. Functional imaging-derived measures were residualized for linear effects of age, mean head framewise displacement, and average network edge weight. Associations between age and brain measures, controlling for modality-specific quality measures, are shown in **Supplemental Figure 1**.

We estimated our bilayer networks using Gaussian Graphical Models (Lauritzen, 1996). Gaussian Graphical Models permit the calculation of weighted partial correlation coefficients, which estimate the conditionally dependent relationships between nodes given the set of all other nodes within the network. For instance, any partial correlation between one node (e.g., family income) and a different node such as the visual system is a statistical relationship that remains after controlling for the associations among these two nodes with all other nodes in the network, regardless of the level of organization (i.e., exposome vs brain network). To estimate all Gaussian Graphical Models, we used the Extended Bayesian Information Criterion graphical least absolute shrinkage and selection operator (EBICglasso, Chen and Chen, 2008; Epskamp et al., 2012; Epskamp and Fried, 2018; Foygel and Drton, 2010; Friedman et al., 2014, 2008; Tibshirani, 1996). The EBICglasso is a regularization method that generates sparser networks by including penalization for more complex models. We set our EBIC tuning parameter to 0.5, considered a high value that prefers simpler, more parsimonious networks (Epskamp and Fried, 2018; Foygel and Drton, 2010). This results in parameter estimates with very small edge weights being set to 0. This procedure reduces the risk of spurious (i.e., false positive) connections within the network.

We calculated Pearson correlations to use as our weighted partial correlation coefficients, and we used pairwise deletion to account for data missingness. For each bilayer network, we estimated two types: one without thresholding and one with thresholding. Thresholding employs a rule for ensuring low false positive rates of edges (Jankova and van de Geer, 2018), which sets edges to zero that are not larger than the threshold in the EBICglasso computation of all considered models, as well as the final returned model. This uses a lower bound for removing the smallest edges, resulting in different edge parameter estimations compared to the unthresholded bilayer network (see **Supplementary Material Figures 2 and 3**). Given the exploratory nature of this study, we did not hypothesize any specific directions (positive or negative) or magnitude (e.g., small or moderate partial correlations) among nodes. All network estimation was done using the packages *bootnet* (version 1.5, Epskamp et al., 2018) and *qgraph* (version 1.9.2, Epskamp et al., 2012).

A common practice in network science is to calculate which nodes have the greatest influence within the network. This property of network nodes is called centrality. Given our interest in inter-network connections between layers (i.e., exposome to brain), we calculated nodal bridge centrality (Jones et al., 2021). Bridge centrality quantifies the extent to which a node (e.g., exposome) forms a partial correlation with another node outside of its community (brain). For our centrality measures, we calculated bridge strength and bridge expected influence separately for each exposome and brain system node (Jones et al., 2021). Bridge strength sums the absolute weights of the partial correlation coefficients for each node to a node from a different community. Bridge expected influence, similar to bridge strength, sums the signed weights of the partial correlation coefficients from each node to another node from a different community. However, an important distinction is that while bridge strength sums the absolute weights, bridge expected influence takes the sign of the partial correlation into account. For example, a node might have two inter-network connections (i.e., partial correlations) to other nodes, one with a magnitude of 1 and the other with a magnitude of -1. In terms of bridge strength centrality, the node would have a bridge strength of 2 while, for bridge expected influence, the node would score 0. To account for differing community sizes between layers (exposome = 14 nodes vs. brain = 7 nodes), we normalized all centrality measures, which divides centrality estimates by their highest possible value, assuming a maximum edge weight of 1. Centrality measures (bridge strength and bridge expected influence) were computed using the *networktools* package (version 1.5.0, Jones, 2022).

We classified nodes as central if their bridge strength and/or bridge expected influence *z*-score was positive and equal to or greater than one standard deviation above the mean. We do not discuss or interpret negative centrality z-score values for our exposome-brain bilayer networks. Central bridge strength nodes were interpreted as nodes that have the most connections, compared to non-central bridge nodes, between network communities. For bridge expected influence, highly central nodes were interpreted as having the greatest relative impact between communities. In all bilayer networks, for both edge weight coefficients and centrality measures, we calculated the correlation stability coefficient, which indicates the maximum proportion of cases that can be dropped from the sample and, with 95% probability, still retain a correlation of 0.7 (i.e., correlation between rank order of centrality in network estimated on full sample with rank order of centrality indices estimated from networks using subsets of the total sample, see (Epskamp and Fried, 2018). We estimated correlation stability coefficients using 2,000 bootstraps and considered a correlation stability value of 0.5 to be stable (Epskamp et al., 2018).

## Results

### Structural Bilayer Networks

Our structural analyses focused on cortical surface area and cortical thickness extracted from a functional network parcellation (Yeo et al., 2011). We present two versions of each network, one that is unthresholded to show a full picture of the possible relationships among nodes, and one that is thresholded, with edges set to zero that are not larger than the threshold both in the EBICglasso computation of all considered models, as well as the final returned model. Bilayer structural networks (**Figure 1**) showed highest clustering within level (intra-network connections), indicating that the exposome variables were highly interrelated, as were brain structure measures. This was especially the case for the unthresholded networks (**Figure 1A-B**), which were denser compared to the thresholded networks (**Figure 1C-D**). The exposome’s neighborhood crime variables (e.g., murder, burglary, robbery, etc.) mostly correlated positively with each other. In the unthresholded networks, the pollution (particulate matter and lead) and socioeconomic status nodes (parent education, family income, percentage of the neighborhood with bachelor’s degrees, neighborhood unemployment) correlated with each other. Again only in the unthresholded networks, particulate matter and blood-lead level mainly connected with neighborhood and family SES variables, while ACEs were associated across every subcluster (neighborhood crime, pollution and SES). Interestingly, in the thresholded bilayer networks, particular matter was ‘kicked out’ of the network, while ACEs either only correlated with blood lead levels (**Figure 1B**) or was also kicked out of the network (**Figure 1D**). Descriptive statistical properties of the structural networks are shown in **Table 3**. Partial correlations among all variables in our exposome-brain structure networks are in **Supplemental Figure 2**.

**Figure 1.**
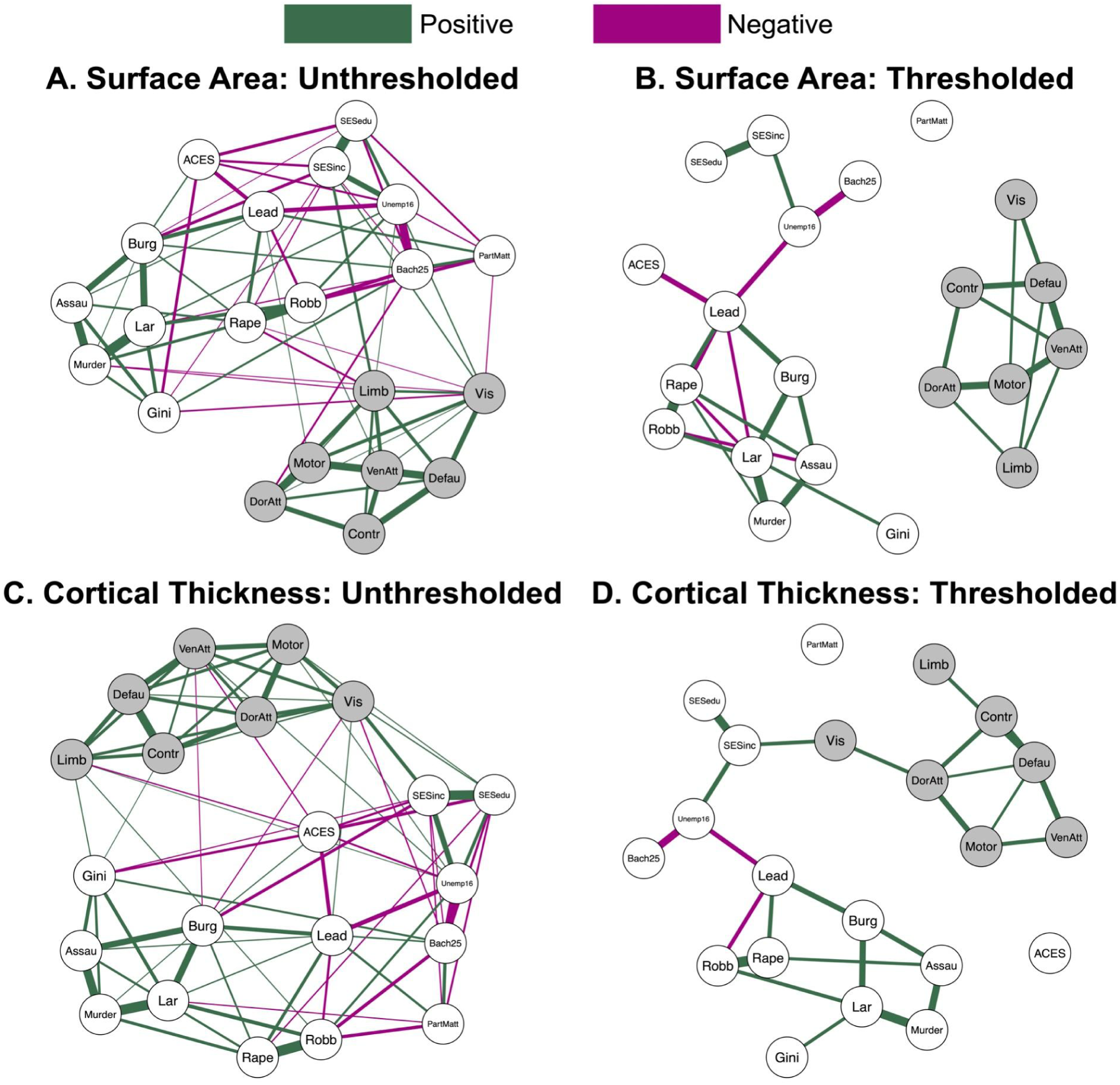
Exposome-Cortical Structure Networks. Multilayer networks were estimated using the Extended Bayesian Information Criterion graphical least absolute shrinkage and selection operator (EBICglasso). We set the tuning parameter to 0.5 and used pairwise deletion to account for data missingness. Partial correlation coefficients (edge weights) were calculated using Pearson method. Green solid lines show positive associations. Magenta lines show negative associations. Thickness of the edge weights indicate the magnitude of the partial correlation (thicker edges show larger partial correlations between nodes). Cortical systems are defined from a seven system parcellation (Yeo et al., 2011): visual (Vis), somatomotor (Motor), dorsal attention (DorAtt), ventral attention (VenAtt), executive control (Contr), default mode (Defau), limbic (Limb). Three exposome variables are reported by parents: income (SESinc), parent education (SESedu), child adverse childhood experiences (ACEs). The other exposome measures are geocoded from census block (see **Supplementary Table 1**): neighborhood incidence of murder (Murder), aggravated assault (Assau), larceny (Lar), rape (Rape), robbery (Robb), burglary (Burg), unemployment over the age of 16 (Unemp16), percent of people over the age of 25 with a Bachelor’s degree or higher (Bach25), Gini index of income inequality (Gini), particulate matter 2.5 (PartMatt), blood lead levels (Lead). A. Surface Area: Unthresholded. B. Surface Area: Thresholded. Edges that are not larger than the threshold of both the EBICglasso computation of all considered models and the final returned model are set to zero. C. Cortical Thickness: Unthresholded. D. Cortical Thickness: Thresholded. Edges that are not larger than the threshold of both the EBICglasso computation of all considered models and the final returned model are set to zero.

**Table 3.** Network properties of the exposome-brain structure (cortical surface area and thickness) bilayer networks.

|  | Exposome-cortical surface area |  | Exposome-cortical thickness |  |
| --- | --- | --- | --- | --- |
|  | Unthresholded | Thresholded | Unthresholded | Thresholded |
| Edge Weight<br>Mean (sd)<br>[range] | .07 (.17)<br>[-.39 – .52] | .18 (.28)<br>[-.45 – .69] | .07 (.15)<br>[-.39 – .51] | .22 (.24)<br>[-.44 – .65] |
| Total Nonzero<br>Edges | 76 | 31 | 78 | 25 |
| Total Negative<br>Edge Number | 26 | 7 | 23 | 3 |
| Nonzero<br>inter-network<br>edges | 13 | 0 | 15 | 1 |
| Negative<br>inter-network<br>edges | 7 | 0 | 5 | 0 |
| Edge weight<br>stability | .67 | .67 | .59 | .59 |
| Bridge strength<br>stability | .28 | N/A | .36 | N/A |
| Bridge expected<br>influence stability | .21 | N/A | .13 | N/A |

We found convergent and divergent patterns in the inter-network (exposome to brain layer) connections in our two structural bilayer networks. In the unthresholded exposome-cortical surface area network (**Figure 1A**), limbic surface area was negatively associated with two crime variables (murder PC = -.02, rape PC = -.07), and positively associated with family income (PC = .10) and neighborhood unemployment (PC = .003). Greater visual surface area was associated with higher parent income (PC = .04) and lower neighborhood income inequality (PC = -.05), but it was also associated with higher neighborhood unemployment (PC = .03), murder index (PC = -.004), and particulate matter concentration (PC = -.01). Surface area of the dorsal attention system was negatively associated with neighborhood education levels (PC = -.06). Surface area of the ventral attention system was positively associated with neighborhood blood lead levels (PC = .01). In the thresholded exposome-cortical surface area network (**Figure 1B**), no between layer connections remained.

For the unthresholded cortical thickness network (**Figure 1C**), greater visual system thickness was associated with higher family income (PC = .14), higher parent education (PC = .03), and higher robbery index (PC = .01). However, visual system thickness showed negative partial correlations with neighborhood education levels (PC = -.03) and burglary index (PC = -.02). Higher ventral attention system thickness was associated with higher neighborhood unemployment (PC = .02), but lower incidence of burglary (PC = -.01) and fewer ACEs (PC = -.03). Limbic system cortical thickness showed partial correlations with the neighborhood incidence of robbery (PC = .02), neighborhood inequality (PC = .02), unemployment in the neighborhood (PC = .004), and ACEs (PC = -.03). Thickness of the somatomotor system correlated positively with parent education (PC = .01) and neighborhood unemployment (PC = .01). Executive control system cortical thickness positively correlated with the Gini Index (PC = .003). For the thresholded cortical thickness network (**Figure 1D**), only one between-layer connection was found, from family income to visual cortical thickness (PC = .20).

We calculated centrality measures only for the unthresholded networks because so few between-layer connections emerged in the thresholded networks. For the exposome-cortical surface area network (**Figure 2A**), four nodes displayed s high bridge strength (*z*-score > 1 SD): parent income (*z* = 2.88), limbic system surface area (*z* = 1.72), rape index (*z* = 1.04), and visual system surface area (*z* = 1.01). Only one node emerged as highly central, for bridge expected influence (**Figure 2B**): parent income (*z* = 3.45). The exposome-cortical thickness bilayer network contained two high-bridge-strength nodes (**Figure 2C**): parent income (*z* = 3.03) and visual system thickness (*z* = 2.32). The same two nodes emerged as highly central in terms of bridge expected influence (**Figure 2D**): family income (*z* = 3.31) and visual system thickness (*z* = 1.45). All centrality estimates were unstable after undergoing bootstrap analysis, revealing that fewer than 50% of the sample could be dropped and maintain a correlation of .70 with the full sample, with .95 probability. However, edge-weights for all networks were stable after bootstrapping. See **Table 3** for estimates of centrality and centrality stability for all exposome-brain structure networks.

**Figure 2.**
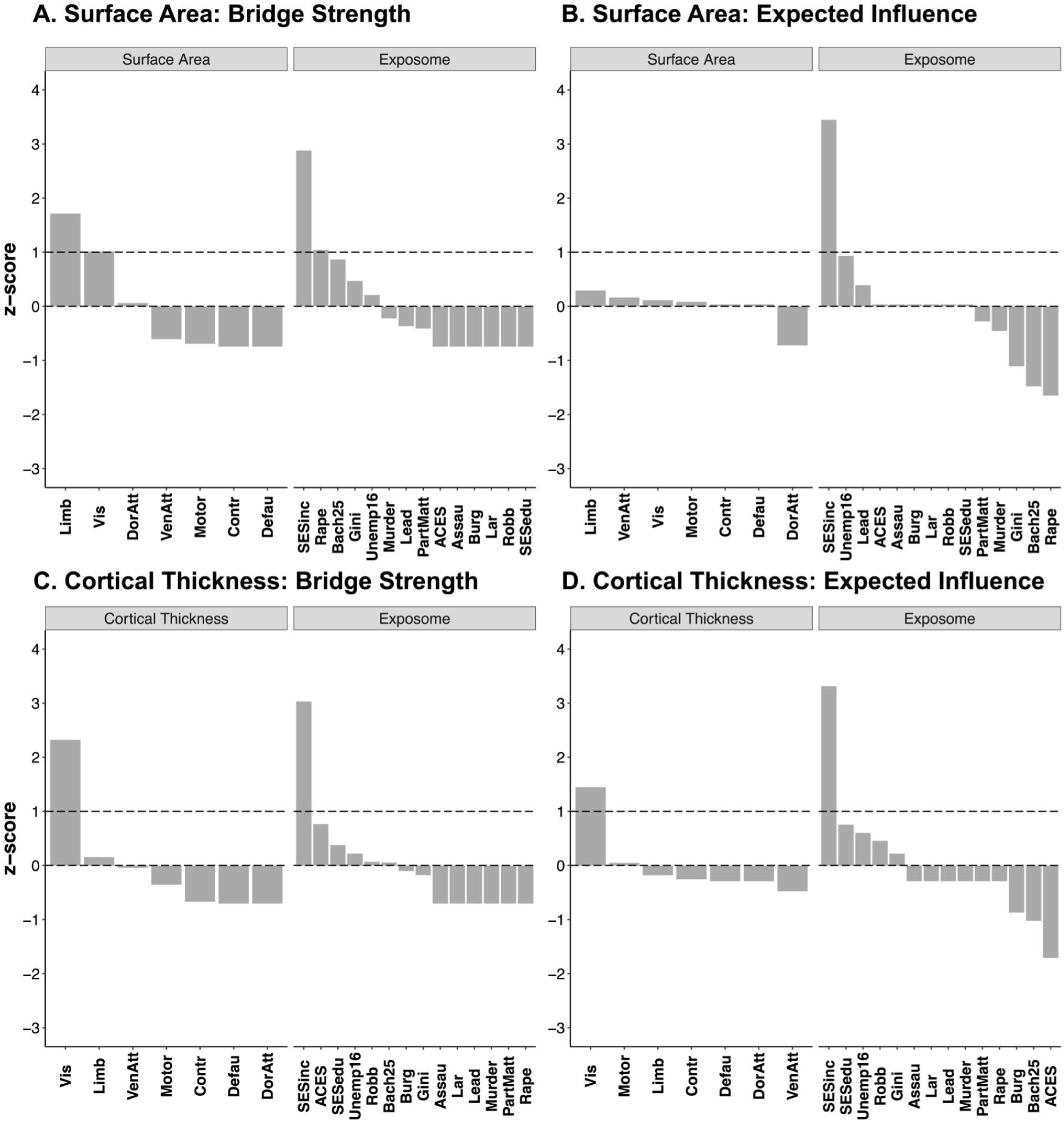
Normalized centrality metrics for the unthresholded exposome-cortical structure networks. A. Surface Area: Bridge Strength. B. Surface Area: Bridge Expected Influence. C. Cortical Thickness: Bridge Strength. D. Cortical Thickness: Bridge Expected Influence.

### Functional Bilayer Networks

We focused on two functional properties of cortex: the participation coefficient and the clustering coefficient, which respectively capture integration and segregation. The participation coefficient was negatively associated with age, and the clustering coefficient was positively associated with age (**Supplemental Figure 3**). As in the structural networks, bilayer functional networks showed strong connectivity among exposome variables and among functional measures. Descriptive statistics of the functional networks are shown in **Table 4**. Partial correlations among all variables in the exposome-brain function networks are in **Supplemental Figure 3**.

**Table 4.**
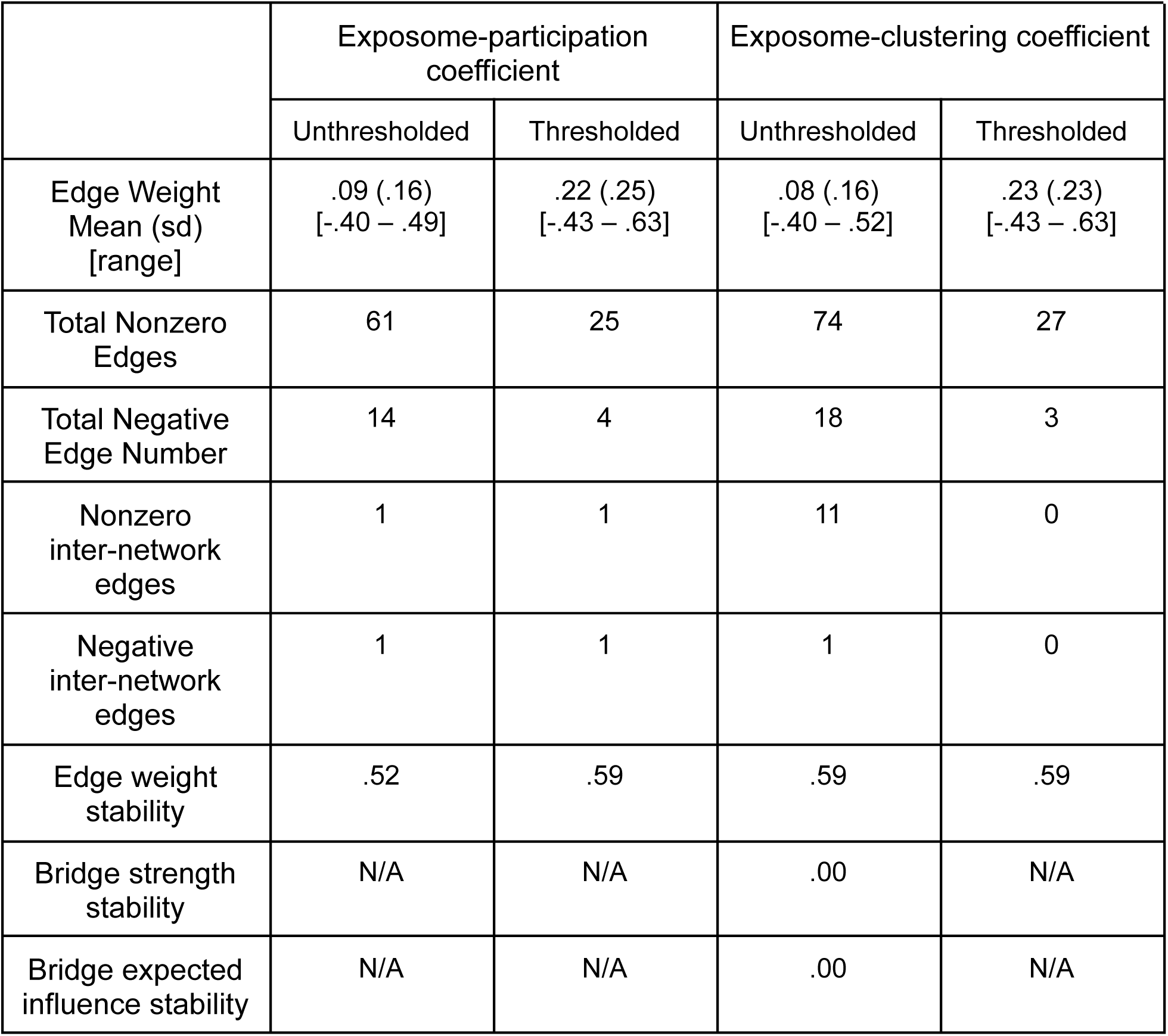
Network properties of the exposome-brain function (participation coefficient and clustering coefficient) bilayer networks.

In the unthresholded exposome-participation coefficient network (**Figure 3A**), lead showed a negative association with the participation coefficient of the dorsal attention system (PC = -.05). In the thresholded exposome-participation coefficient network (**Figure 3B**), lead negatively correlated with the participation coefficient of the default mode system (PC = -.20). In the unthresholded exposome-clustering coefficient network (**Figure 3C**), lead showed a positive association with the clustering coefficient of the default mode system (PC = .04). Particulate matter was positively associated with the clustering coefficient of the control system (PC = .02). The visual system positively correlated with employment (PC = .02), while the motor system negatively correlated with family income (PC = -.02). Family education positively correlated with the default mode system (PC = .01). Lastly, the dorsal attention system showed positive associations with inequality (PC = .01) and larceny (PC = .01). There were several other between-layer connections, but all had a partial correlation below .005 (see **Supplemental Figure 3C**). In the thresholded exposome-clustering coefficient network (**Figure 3D**), no between-layer edges emerged.

**Figure 3.**
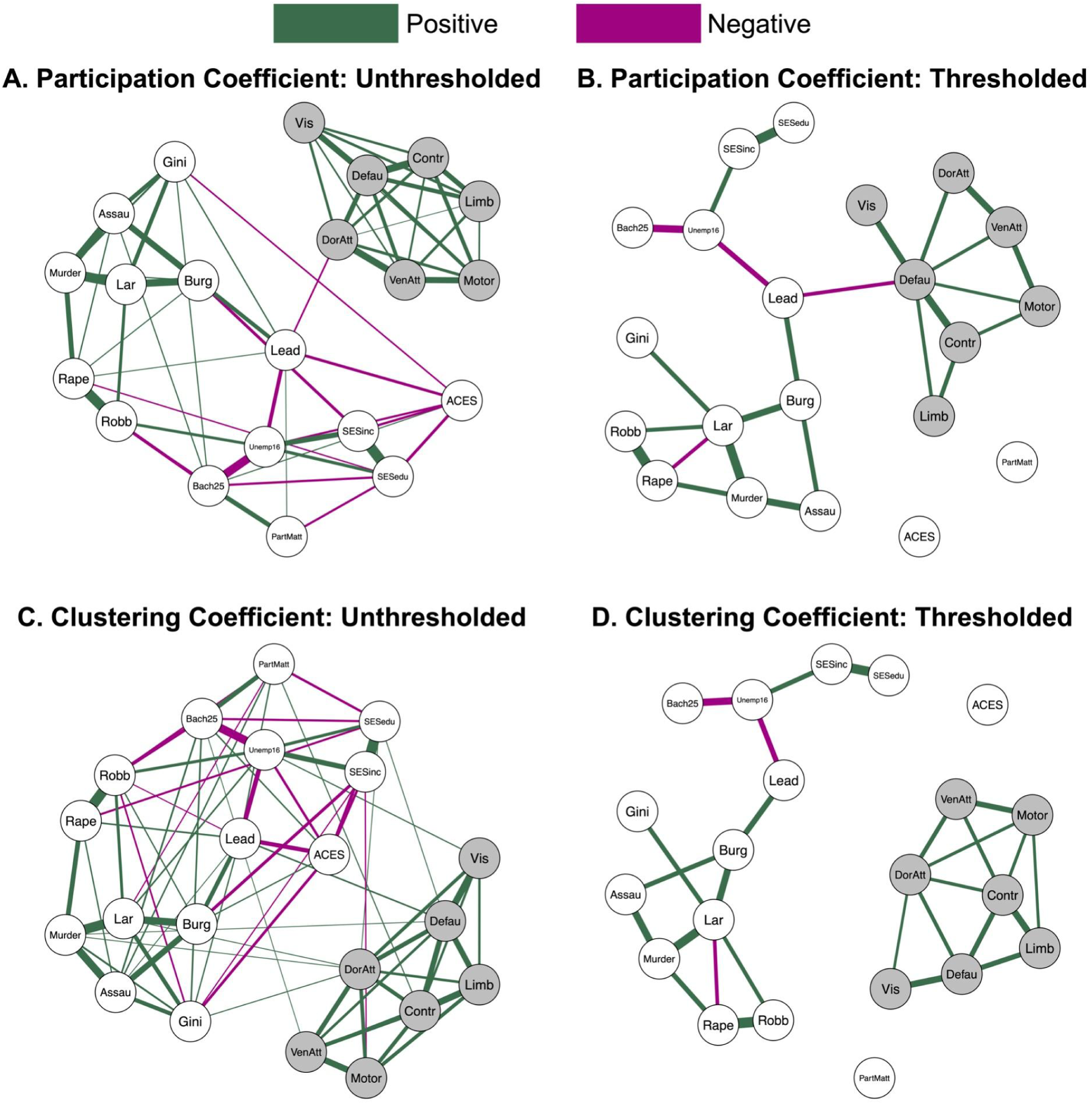
Exposome-Cortical Function Networks. Multilayer networks were estimated using the Extended Bayesian Information Criterion graphical least absolute shrinkage and selection operator (EBICglasso). We set the tuning parameter to 0.5 and used pairwise deletion to account for data missingness. Partial correlation coefficients (edge weights) were calculated using Pearson method. Green solid lines show positive associations. Magenta lines show negative associations. Thickness of the edge weights indicate the magnitude of the partial correlation (thicker edges show larger partial correlations between nodes). Cortical systems are defined from a seven system parcellation (Yeo et al., 2011): visual (Vis), somatomotor (Motor), dorsal attention (DorAtt), ventral attention (VenAtt), executive control (Contr), default mode (Defau), limbic (Limb). Three exposome variables are reported by parents: income (SESinc), parent education (SESedu), child adverse childhood experiences (ACEs). The other exposome measures are geocoded from census block (see **Supplementary Table 1**): neighborhood incidence of murder (Murder), aggravated assault (Assau), larceny (Lar), rape (Rape), robbery (Robb), burglary (Burg), unemployment over the age of 16 (Unemp16), percent of people over the age of 25 with a Bachelor’s degree or higher (Bach25), Gini index of income inequality (Gini), particulate matter 2.5 (PartMatt), blood lead levels (Lead). A. Participation Coefficient: Unthresholded. B. Participation Coefficient: Thresholded. Edges that are not larger than the threshold of both the EBICglasso computation of all considered models and the final returned model are set to zero. C. Clustering Coefficient: Unthresholded. D. Clustering Coefficient: Thresholded. Edges that are not larger than the threshold of both the EBICglasso computation of all considered models and the final returned model are set to zero.

We only calculated bridge centrality for the unthresholded exposome-clustering coefficient network (**Figure 3C**) due to the sparse or nonexistent bilayer connections for the functional networks. For the exposome-clustering coefficient network (**Figure 4A**), three nodes displayed high bridge strength (*z*-score > 1 SD): incidence of high blood lead levels (*z* = 2.88), default mode network clustering coefficient (*z* = 1.41), and parent income (*z* = 1.20). Two nodes emerged as highly central, for bridge expected influence (**Figure 4B**): incidence of high blood lead levels (*z* = 2.61), default mode network clustering coefficient (*z* = 1.40), and neighborhood rate of unemployment (*z* = 1.05). See **Table 4** for edge weight stability estimates, descriptive statistics for all exposome-brain function networks, and the bridge centrality stability estimates for the exposome-clustering coefficient network..

**Figure 4.**
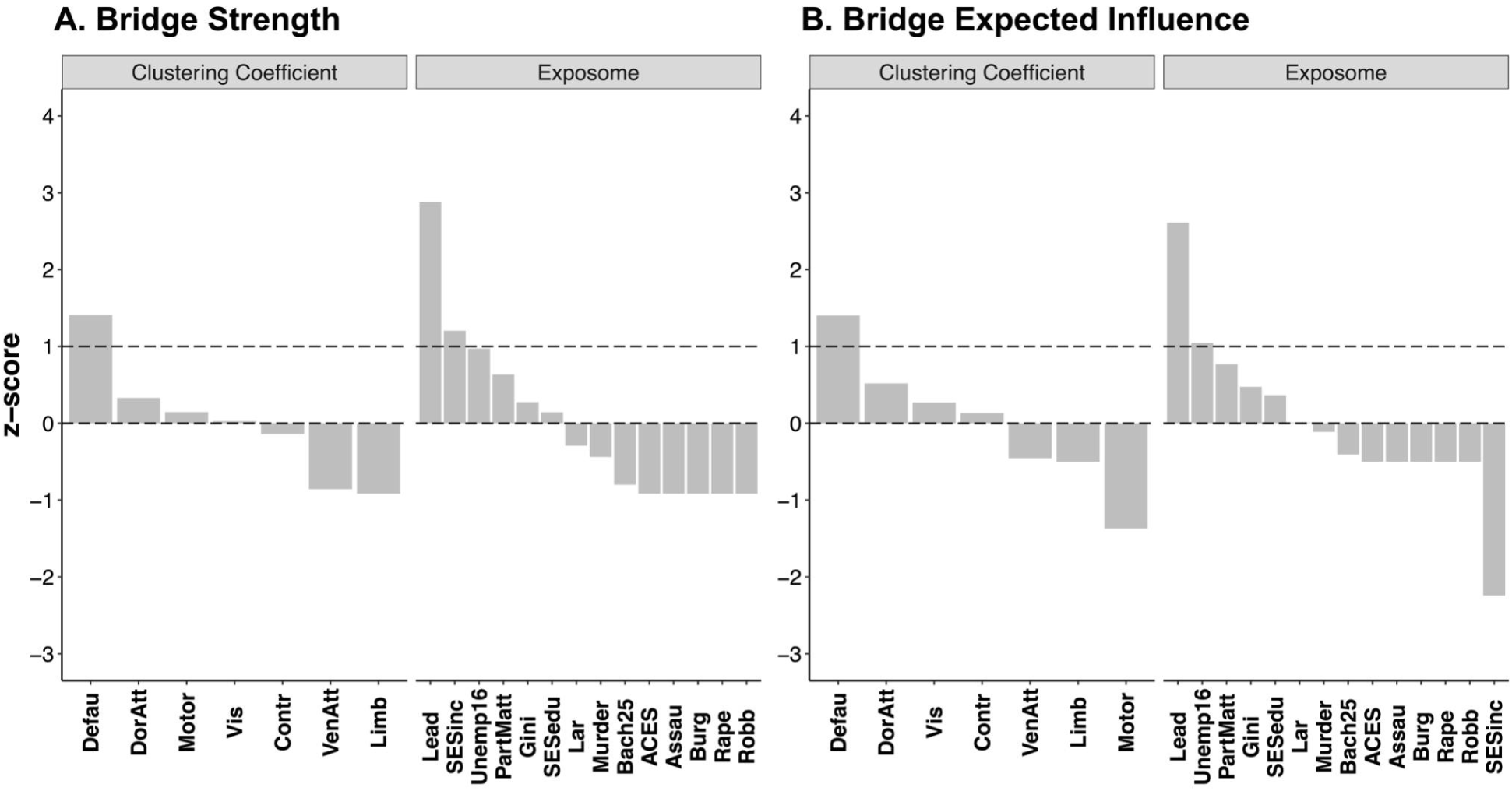
Normalized centrality metrics for the unthresholded exposome-clustering coefficient network. A. Bridge Strength. B. Bridge Expected Influence.

## Discussion

In this study, we built exposomes connecting family socioeconomic status and child adversity with neighborhood measures of socioeconomic status, crime, and pollution. We linked the exposomes to measures of cortical structural and functional maturation. This approach yielded three key results. First, lower family income was associated with thinner visual cortex. Second, greater neighborhood incidence of murder was associated with less limbic surface area. Third, greater neighborhood incidence of high blood lead levels was associated with lower participation coefficient and clustering coefficient in the default mode system. Taken together, these results demonstrate the potential of multilayer network models to generate new insights into how children’s experiences influence their brain development. These models might make previously unknown connections apparent as they take into account all possible connections in the multilayer network (e.g., highly correlated environmental exposures and brain measures).

Our multilayer network models also uncovered associations previously found in studies which focused on single exposures. For example, lower family income was associated with thinner visual cortex. A meta-analysis of associations between SES and brain structure across the first two decades of life found associations in visual areas, but these associations were not specific (Rakesh and Whittle, 2021). In contrast, an analysis correlating income with cortical thickness in early childhood (ages 4-7) found a specific relationship in visual cortex, with no other areas showing a significant relationship (Leonard et al., 2019). If the stress associated with low financial resources accelerates brain development, the association between SES and cortical thinning might be strongest in early developing regions early in childhood. Indeed, in this dataset, visual cortical thickness strongly correlates with age while other systems may not yet be thinning. It is also possible that the relationship between income and visual cortical thickness is driven by differences in visual experience, for example differences in exposure to novel or complex objects or differences in how caregivers guide visual attention (Rosen et al., 2019; Werchan et al., 2019). Other explanations for the strength of the visual effects are possible. Visual cortex is also closer to head coil elements than other areas, especially for smaller heads, and may be less susceptible to movement artifacts because children rest on the back of their heads in the scanner (Alexander-Bloch et al., 2016; Rosen et al., 2018).

The finding that neighborhood violence shows a specific association with limbic structure is consistent with the hypothesis that threats have a specific influence on limbic areas of the brain (De Brito et al., 2013; Edmiston, 2011; Hanson et al., 2010; McLaughlin et al., 2014). Neighborhood violence has been repeatedly linked to children’s cognition and behavior, even in young children (McCoy et al., 2023; Sharkey, 2010). For example, a study in Brazil showed that recent neighborhood violence was associated with reduced performance on cognitive tests in 3-year-old children (McCoy et al., 2023). Other work using a geocoded measure of murder found that neighborhood murder was associated with inflammatory activity, but only in children with low resting-state functional connectivity of the salience or central executive network, suggesting that children differ in their sensitivity to neighborhood crime (Miller et al., 2021). This work highlights the need for further study of how young children are influenced by neighborhood violence to investigate possible mechanisms of how and in which environments (proximal or distal) it affects brain development.

In the functional-exposome networks, default mode network (DMN) measures were associated with neighborhood incidence of high blood lead levels in children. Lead exposure has been linked to lower cortical volume and surface area in 9-10-year-olds (Marshall et al., 2020). Childhood lead exposure has also been shown to relate to lower cortical surface area in midlife (Reuben et al., 2020). The literature on lead effects on functional networks is limited, but there are studies on other toxins. For example, pesticide exposure has been linked to greater clustering coefficient in the DMN of 8-year-olds (Bahrami et al., 2022). Vehicle exhaust exposure has been linked to weaker functional connectivity between medial frontal and angular gyrus hubs of the DMN in 8-12-year-olds (Pujol et al., 2016). In our study, higher neighborhood lead exposure was associated with lower DMN participation coefficient. Because the participation coefficient is negatively associated with age as network architecture is refined, this result is consistent with the possibility of accelerated functional maturation (Fair et al., 2009, 2007; Tooley et al., 2022b, 2020). DMN effects may reflect the network’s proximity to major arteries and venous drainage (Satterthwaite et al., 2014). In animal models, it has been shown that a variety of toxins are first deposited along the midline of the brain near major arteries (Kesler et al., 2013; Mills et al., 2016). It is also possible that lead exposure is associated with inflammatory responses, which is consistent with a documented relationship between interleukin-6 levels in blood and functional connectivity of the DMN in adults (Marsland et al., 2017).

This study has a number of limitations. Because of the small sample size, we constrained the number of environmental and brain features in the models. Our sample and geocoded data did not include measures on some key dimensions of early experiences, in particular, positive and protective experiences, and on the duration and timing of exposures. Our geocoded data can tell us about average experiences in a census block but do not provide individual-level experiences with crime or exposure to environmental toxins, which could differ along many aspects of family identities, including race, gender, and age. Likewise, the measures used here were not all raw measures of exposures; the Market Profile crime indices and particulate matter concentration in a census block are predictions from other available county or city-level data. We also focused only on the cortex, dividing it into relatively large regions. Future work with larger samples could expand the exposome, and facilitate a more granular investigation of the brain, including subcortical gray matter and white matter. We did not have the power to look at age by exposome interactions on brain development to ask whether the patterns we see look different in younger vs. older children. This is further evidenced in the low stability of all our bilayer networks, which suggests that the relationships observed in this study have limited robustness. Larger longitudinal datasets are necessary to better evaluate developmental timing hypotheses. Even in the constrained models, choices about included variables, thresholding, sample size, and/or decisions about how to deal with data missingness (e.g., data imputation to protect against estimation bias, see Liu et al., 2023) can change network layouts, so we see the specific network architecture as a description of the data in this sample under a set of sensible methodological choices rather than definitive evidence for specific associations among exposome and brain layers. Despite these limitations, recent work has demonstrated that with sample sizes of ∼300, the EBICglasso estimator performs well at recovering a network structure resembling the true network and the strongest edges (Isvoranu and Epskamp, 2021). Whether this finding translates to our smaller sample size remains unknown. Moreover, the EBICglasso estimator has shown good sensitivity regarding the detection of bridge edges, especially those strongest within the network. However, this comes at the expense of lower precision and specificity, which suggests that some bridge edges might be false positives. Altogether, this points to the need for future research to expand upon our findings using larger sample sizes (e.g., the ABCD study, see Casey et al., 2018) and comparisons among different estimation methods. However, the current sample benefits from containing detailed exposure measures and neuroimaging data from a community sample of children from diverse backgrounds (see demographic characteristics from **Table 1**), which makes it unique to explore these associations.

To conclude, we took a multilayer network approach to characterize how a diverse set of early experiences in children’s homes and neighborhoods relate to cortical structure and function. This approach has the potential to inform not only basic science theories of brain development, but also intervention approaches. Our specific results suggest that family income is central, connecting multiple types of adversity to children’s brain structure. Our results also suggest that lead exposure could be impacting default mode network function above and beyond stress exposure. If these results replicate, they could be seen as additional justification for policies like unconditional cash transfers and lead monitoring and abatement, though the empirical support for these policies is already extensive (for review, see Blair and Raver, 2016; Farah, 2018). Repeating these analyses in different geographic contexts could test how regional differences in the coincidence of exposures (e.g., spatial correlations of crime and pollution), and differences in policies to mitigate the impact of poverty (Weissman et al., 2023) and lead exposure, change associations with child brain development. Experimental changes in nodes of these networks, like cash transfers (Baby’s First Years; Noble et al., 2021), will give us even stronger insights into the validity of the networks we describe here. Together, this study highlights the utility of connecting explanatory levels by applying a multilayer approach to not only replicate previous findings but also provide new directions for further understanding the complex interactions between childhood brain development and their environments.

## Code And Data Availability Statement

All code for this study is available on Open Science Framework: https://osf.io/sfymz/. Due to the sensitive nature of the geocoded measures, the data files for this study are available with submission of evidence of human subjects training.

## Acknowledgments

We thank the families who participated in this research. This study was supported by the Jacobs Foundation Early Career Award (A.P.M.), the National Institute on Drug Abuse (Award 1R34DA050297-01 to A.P.M.), the University of Pennsylvania’s Data Driven Discovery Data Science Postdoctoral Fellowship (I.L.S.), and the National Science Foundation (Graduate Research Fellowships to A.L.B., C.L.M., and U.A.T.). This work was also supported by an Azrieli Global Scholar award to A.P.M. from the Canadian Institute for Advanced Research (CIFAR), and an NSF CAREER award to A.P.M. (DRL 2045095). We also thank Stephanie Bugden, Jasmine Forde, Katrina Simon, Sophie Sharp, Isis Cowan, Maayan Ziv, Yoojin Hahn, and the Changing Brain Lab for their help with data acquisition.

## SUPPLEMENTARY MATERIAL

**Supplemental Table 1.** Demographic and geocoded information of participants in the structural neuroimaging sample and the participants who participated at their first scan session but were excluded. *Survey form allowed parents to endorse more than one racial identity hence the sum of racial identities is greater than 100%. ^十^Market Profile variables indices represent deviations from the U.S. average rate of that specific crime - U.S. mean rate of a specific crime is 100. Comparisons between continuous variables, such as crime indices, Gini index, age, income, years of education, PM2.5 concentrations, and others were performed using Wilcoxon rank sum t-tests; categorical variables (specific race categories, ethnicity, and sex) were compared with Fisher exact t-tests. Two variables were found to be significantly different (*p < 0.05*) between the two samples: burglary index and age.

|  | Structural imaging<br>(*n* = 170) | Excluded<br>participants<br>(*n* = 42) | Comparison test<br>p-value |
| --- | --- | --- | --- |
| Age (years, mean [SD] {range}) | 6.34 [1.40]<br>{4.05-10.59} | 5.66 [0.95]<br>{4.18-7.52} | **0.003\*\*** |
| Sex (*n* [%]) | 89 [52.4%] female<br>81 [47.6%] male<br>0 [0%] other | 20 [47.6%] female<br>22 [52.4%] male<br>0 [0%] other | 0.6 |
| Race (%) | 50.6 % Black<br>41.8 % White<br>11.8 % Asian<br>1.8 % Native<br>Hawaiian or Other<br>Pacific Islander<br>1.8% American<br>Indian or Alaska<br>Native<br>6.5% Other | 47.6 % Black<br>40.5 % White<br>11.9 % Asian<br>2.4 % Native<br>Hawaiian or Other<br>Pacific Islander<br>2.4% American<br>Indian or Alaska<br>Native<br>0.0% Other | (0.9) Black<br>(>0.9) White<br>(>0.9) Asian<br>(>0.9) Native<br>Hawaiian or Other<br>Pacific Islander<br>(>0.9) American<br>Indian or Alaska<br>Native<br>(0.13) Other |
| Ethnicity (%) | 11.3%<br>Latinx/Hispanic | 16.7%<br>Latinx/Hispanic | 0.4 |
| Family Income (thousands of \$,<br>mean [SD]) | 80.1 [68.2] | 92.4 [63.4] | 0.4 |
| Parent Education (years, mean<br>[SD]) | 14.88 [2.72] | 15.26 [2.79] | 0.4 |
| Adverse Childhood Experiences<br>(ACEs)<br>(mean, [SD]) | 0.97 [1.35] | 1.12 [1.31] | 0.2 |
| Neighborhood unemployment, (%,<br>mean [SD]) | 34.7 [24.5] | 36.3 [27.9] | >0.9 |
| Percentage of people (25+ years<br>old) with a Bachelor's degree in<br>the neighborhood (% , mean [SD]) | 8.3 [5.4] | 9.3 [6.7] | 0.6 |
| Percentage incidence of high<br>blood levels in neighborhood (% ,<br>mean [SD]) | 6.3 [3.4] | 6.1 [2.6] | 0.7 |
| Average particulate matter<br>concentration in neighborhood<br>(µg/m<sup>3</sup>, mean [SD]) | 1.30 [0.33] | 1.29 [0.33] | >0.9 |
| Murder Index<sup>+</sup> (unitless, mean [SD]) | 110.5 [50.8] | 92.5 [50.5] | 0.11 |
| Rape Index<sup>+</sup> (unitless, mean [SD]) | 94.0 [53.0] | 81.7 [50.3] | 0.2 |
| Robbery Index<sup>+</sup> (unitless, mean [SD]) | 90.1 [58.2] | 79.6 [56.8] | 0.3 |
| Assault Index<sup>+</sup> (unitless, mean [SD]) | 113.2 [41.8] | 92.4 [53.2] | 0.054 |
| Larceny Index<sup>+</sup> (unitless, mean [SD]) | 115.2 [41.8] | 103.0 [45.0] | 0.2 |
| Burglary Index<sup>+</sup> (unitless, mean [SD]) | 112.1 [50.4] | 88.4 [56.0] | **0.029\*** |
| Gini Index (unitless, mean [SD]) | 0.460 [0.065] | 0.454 [0.057] | 0.9 |

**Supplemental Table 2.** Demographic and geocoded information of participants in the functional neuroimaging sample and the earliest available scan session of the participants who participated in at least one scan session (including those with an earliest scan session after the first) but were excluded. *Survey form allowed parents to endorse more than one racial identity hence the sum of racial identities is greater than 100%. ^十^Market Profile variables indices represent deviations from the U.S. average rate of that specific crime - U.S. mean rate of a specific crime is 100. Comparisons between continuous variables, such as crime indices, Gini index, age, income, years of education, PM2.5 concentrations, and others were performed using Wilcoxon rank sum t-tests; categorical variables (specific race categories, ethnicity, and sex) were compared with Fisher exact t-tests. One variable was found to be significantly different (*p < 0.05*) between the two samples: age.

| | Functional imaging sample<br>($n = 130$) | First available session for excluded participants<br>($n = 87$) | Comparison test p-value |
| --- | --- | --- | --- |
| Age (years, mean [SD] {range}) | 6.64 [1.35]<br>{4.11-10.59} | 5.73 [1.21]<br>{4.05-9.98} | **<0.001\*\*\*** |
| Sex (n [%]) | 69 [53.1%] female<br>61 [46.9%] male<br>0 [0%] other | 42 [48.3%] female<br>45 [51.7%] male<br>0 [0%] other | 0.5 |
| Race* (%) | 51.5 % Black<br>42.3 % White<br>13.8 % Asian<br>2.3% Native<br>Hawaiian or Other<br>Pacific Islander<br>1.5% American<br>Indian or Alaska<br>Native<br>3.1% Other | 48.3 % Black<br>40.2 % White<br>9.2 % Asian<br>0.0% Native<br>Hawaiian or Other<br>Pacific Islander<br>2.3% American<br>Indian or Alaska<br>Native<br>8.0% Other | (0.7) Black<br>(0.8) White<br>(0.4) Asian<br>(0.3) Native<br>Hawaiian or Other<br>Pacific Islander<br>(>0.9) American<br>Indian or Alaska<br>Native<br>(0.12) Other |
| Ethnicity (%) | 10.9%<br>Latinx/Hispanic | 14.0%<br>Latinx/Hispanic | 0.5 |
| Family Income (thousands of \$, mean [SD]) | 85.7 [68.2] | 75.7 [65.8] | 0.3 |
| Parent Education (years, mean [SD]) | 15.09 [2.72] | 14.76 [2.72] | 0.4 |
| Adverse Childhood Experiences (ACEs) (mean, [SD]) | 1.02 [1.31] | 0.99 [1.45] | 0.7 |
| Neighborhood unemployment, (% mean [SD]) | 34.3 [23.9] | 35.2 [26.6] | >0.9 |
| Percentage of people (25+ years old) with a Bachelor's degree in the neighborhood (% mean [SD]) | 8.3 [5.4] | 9.1 [6.1] | 0.4 |
| Percentage incidence of high blood levels in neighborhood (% mean [SD]) | 6.1 [3.4] | 6.8 [3.1] | 0.2 |
| Average particulate matter concentration in neighborhood ($\mu\text{g}/\text{m}^3$, mean [SD]) | 1.32 [0.33] | 1.29 [0.33] | 0.4 |
| Murder Index<sup>+</sup> (unitless, mean [SD]) | 108.3 [49.2] | 105.8 [53.1] | 0.8 |
| Rape Index<sup>+</sup> (unitless, mean [SD]) | 91.5 [52.2] | 92.5 [53.5] | >0.9 |
| Robbery Index<sup>+</sup> (unitless, mean [SD]) | 88.4 [58.0] | 88.5 [58.4] | >0.9 |
| Assault Index<sup>+</sup> (unitless, mean [SD]) | 109.3 [45.0] | 111.5 [52.9] | 0.5 |
| Larceny Index<sup>+</sup> (unitless, mean [SD]) | 111.7 [42.2] | 115.3 [42.2] | 0.4 |
| Burglary Index<sup>+</sup> (unitless, mean [SD]) | 108.4 [52.2] | 108.7 [51.9] | >0.9 |
| Gini Index (unitless, mean [SD]) | 0.458 [0.063] | 0.461 [0.066] | 0.7 |

**Supplemental Table 3.** Geocoded exposome variables.

| Measure | Definition | Source |
| --- | --- | --- |
| Murder Index | Neighborhood incidence of murder, as defined as the act of one person taking another person's life outside of the categories of negligence, suicide, and accident | Market Profile |
| Aggravated Assault Index | Neighborhood incidence of violent attacks with the attempt to inflict serious bodily harm and/or potentially death | Market Profile |
| Larceny Index | Neighborhood incidence of theft of personal property of varying value without force or fraud and can be thus classified as a non-violent crime | Market Profile |
| Rape Index | Neighborhood incidence of sexual assault, attempts, or other kinds of assault with the intent to sexually assault, excluding statutory rape and incest | Market Profile |
| Robbery Index | Neighborhood incidence of taking something of value, care, or custody using violence, force, and/or threat of force | Market Profile |
| Burglary Index | Neighborhood incidence of unlawful entry (or attempt) into a structure to commit a felony or theft. It does not by definition require force to enter | Market Profile |
| Unemployment over the age of 16 | Neighborhood proportion of people older than 16 years old in the labor force that are not employed, not including people such as stay-at-home parents, students, incarcerated or otherwise institutionalized individuals, and retired or seasonal workers | American Community Survey |
| People over 25 with a Bachelor's degree or higher | Neighborhood proportion of people over the age of 25 with some form of higher education qualification (Bachelor's, Master's, professional degree (MD, JD), or PhD) | American Community Survey |
| Gini Index | Neighborhood income inequality index, operationalized by taking the difference between the actual income distribution curve and the perfect equality curve. The higher the index, the higher the inequality | American Community Survey |
| Particulate matter 2.5 (PM2.5) | Neighborhood concentration estimate of fine inhalable particles greater than 2.5 $\mu\text{m}$ in diameter such as industrial pollutants, exhaust fumes, among other harmful inhalants | Environmental Protection Agency |
| Blood lead levels | Neighborhood percentage of children with blood lead levels above 5 $\mu\text{g/mL}$ | Philadelphia Department of Public Health |

**Supplemental Figure 1.**
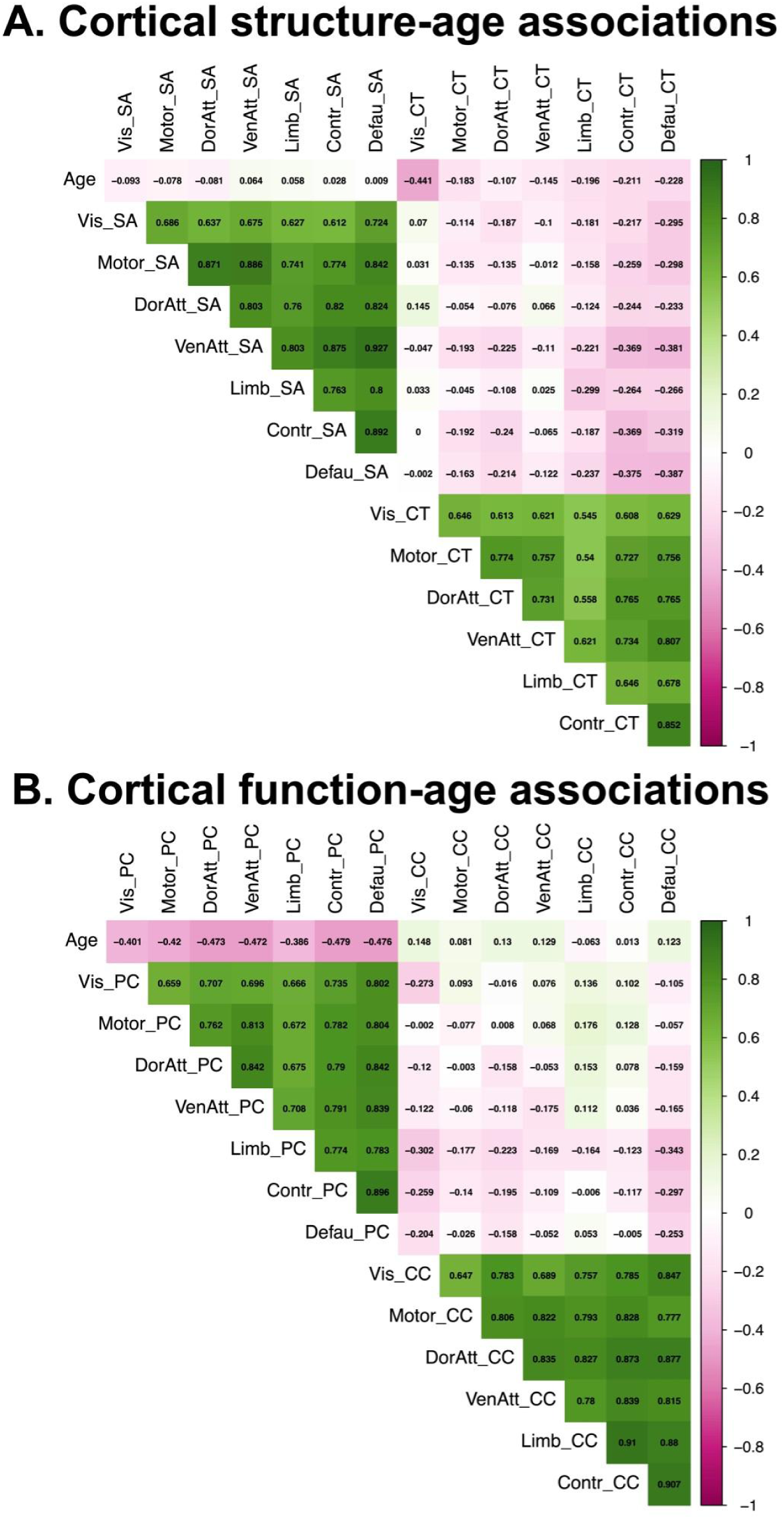
Associations between age and brain measures. A. Cortical Structure measures. Measures include surface area (SA) and cortical thickness (CT). Structural metrics were residualized for image quality rating. B. Cortical Function Measures. Measures include participation coefficient (PC) and clustering coefficient (CC). Functional metrics were residualized for mean head framewise displacement and average network non-zero edge-weight. Cortical systems are defined from a seven system parcellation (Yeo et al., 2011): visual (Vis), somatomotor (Motor), dorsal attention (DorAtt), ventral attention (VenAtt), executive control (Contr), default mode (Defau), limbic (Limb).

**Supplemental Figure 2.**
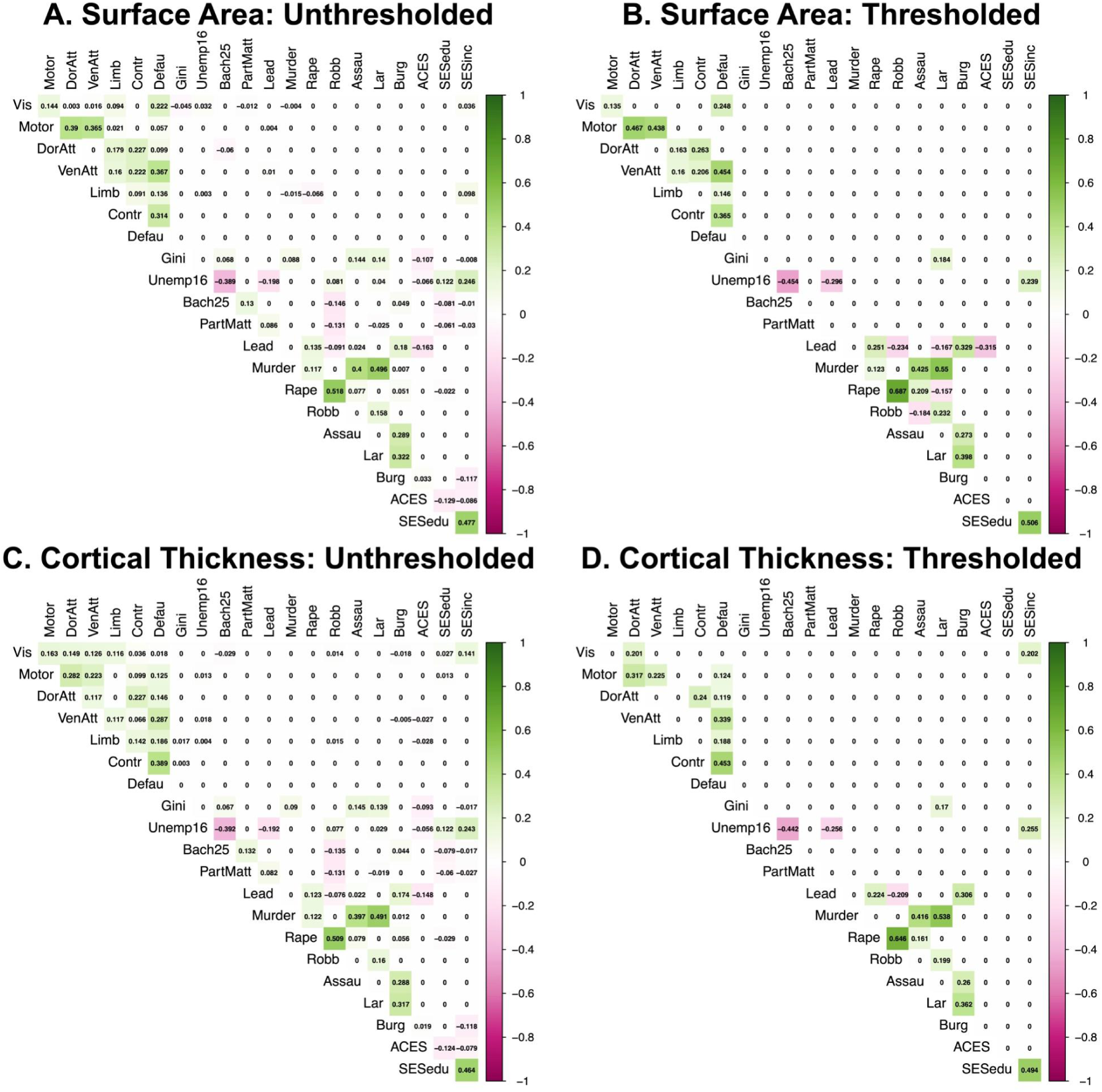
Exposome-Cortical Structure Partial Correlations. Positive partial correlations are shown in green, and negative partial correlations are shown in magenta. Cortical systems are defined from a seven system parcellation (Yeo et al., 2011): visual (Vis), somatomotor (Motor), dorsal attention (DorAtt), ventral attention (VenAtt), executive control (Contr), default mode (Defau), limbic (Limb). Three exposome variables are reported by parents: income (SESinc), parent education (SESedu), child adverse childhood experiences (ACEs). The other exposome measures are geocoded from census block (see Supplementary Table 1): neighborhood incidence of murder (Murder), aggravated assault (Assau), larceny (Lar), rape (Rape), robbery (Robb), burglary (Burg), unemployment over the age of 16 (Unemp16), percent of people over the age of 25 with a Bachelor’s degree or higher (Bach25), Gini index of income inequality (Gini), particulate matter 2.5 (PartMatt), blood lead levels (Lead). A. Surface Area: Unthresholded. B. Surface Area: Thresholded. Edges that are not larger than the threshold of both the EBICglasso computation of all considered models and the final returned model are set to zero. C. Cortical Thickness: Unthresholded. D. Cortical Thickness: Thresholded. Edges that are not larger than the threshold of both the EBICglasso computation of all considered models and the final returned model are set to zero.

**Supplemental Figure 3.**
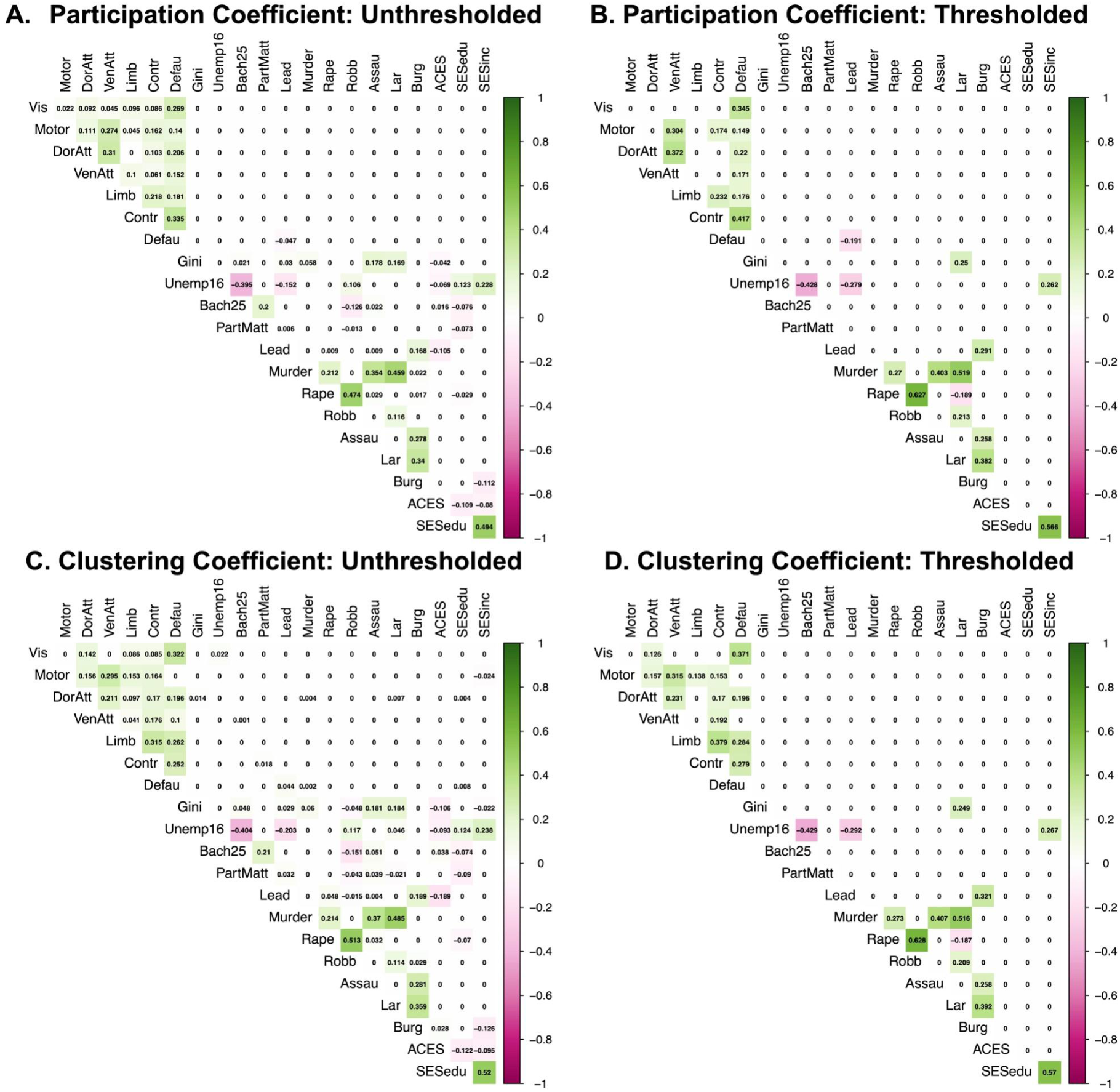
Exposome-Cortical Function Partial Correlations. Positive partial correlations are shown in green, and negative partial correlations are shown in magenta. Cortical systems are defined from a seven system parcellation (Yeo et al., 2011): visual (Vis), somatomotor (Motor), dorsal attention (DorAtt), ventral attention (VenAtt), executive control (Contr), default mode (Defau), limbic (Limb). Three exposome variables are reported by parents: income (SESinc), parent education (SESedu), child adverse childhood experiences (ACEs). The other exposome measures are geocoded from census block (see Supplementary Table 1): neighborhood incidence of murder (Murder), aggravated assault (Assau), larceny (Lar), rape (Rape), robbery (Robb), burglary (Burg), unemployment over the age of 16 (Unemp16), percent of people over the age of 25 with a Bachelor’s degree or higher (Bach25), Gini index of income inequality (Gini), particulate matter 2.5 (PartMatt), blood lead levels (Lead).). A. Participation Coefficient: Unthresholded. B. Participation Coefficient: Thresholded. Edges that are not larger than the threshold of both the EBICglasso computation of all considered models and the final returned model are set to zero. C. Clustering Coefficient: Unthresholded. D. Clustering Coefficient: Thresholded. Edges that are not larger than the threshold of both the EBICglasso computation of all considered models and the final returned model are set to zero.

## Supplemental Methods

### Evaluating a sum score approach to modeling neighborhood crime

Sum scores are commonly used as an aggregate metric of a particular measure of interest (e.g., IQ, SES, etc.). We considered creating a sum score of total neighborhood crime composed of our murder, aggravated assault, larceny, rape, robbery, and burglary crime indices. However, sum scores often misrepresent the underlying construct they are attempting to capture (e.g., clinical diagnoses) if the assumptions underlying the creation of the sum score are not met (see McNeish and Wolf, 2020). For example, we could assume that all neighborhood crime indices are related equally to a latent construct (e.g., Neighborhood Crime), regardless of the nature of the crime (e.g., larceny vs aggravated assault). In this instance, we should constrain the weights of all neighborhood crime indices to be identical when creating the total crime sum score. However, this assumption might be false if any of the crime indices differentially relate to the Neighborhood Crime latent construct (i.e., have non-identical factor loading estimates). In this case, it would instead be appropriate to assign the weight of each crime index depending on its relation to the Neighborhood Crime latent construct.

Furthermore, we might not be justified to use only one sum score to capture neighborhood crime. Instead, neighborhood crime might be better characterized using two or more latent constructs. For example, we can group the individual crime indices into two categories: nonviolent (larceny, robbery and burglary) and violent (rape, aggravated assault and murder). We classify robbery as a nonviolent crime since its definition includes the possibility of stealing an item without physical force (e.g., only the threat of force). Establishing whether neighborhood crime is better represented as a single or multi-factor construct has important implications. For example, if the covariance structure among the crime indices is better represented by two (Non-Violent Crime and Violent Crime) latent constructs rather than one (Neighborhood Crime), it suggests that neighborhood crime is multidimensional, with the potential that these two categories of neighborhood crimes have distinct effects on childhood structural and functional brain development.

Structural equation modeling, used for sum score analyses of our exposome neighborhood crime data (aggravated assault, burglary, larceny, murder, rape, and robbery), was completed using the lavaan (version 0.6-12; Rosseel, 2012) and semPlot (version 1.1.5; Epskamp, 2022) packages for factor model estimation and visualization, respectively. Structural equation modeling (SEM) is a multivariate statistical modeling approach that calculates the correlations (factor loadings) between observed (e.g., neighborhood crime indices) and latent variables (e.g., Neighborhood Crime) (Beaujean, 2014). Moreover, SEM permits the estimation of the fit of the theorized model compared to a baseline model to help determine how well your model captures the covariance structure of your data. We used the robust full information maximum likelihood estimator (FIML) with a Yuan-Bentler scaled test statistic (MLR) and robust standard errors (Schermelleh-Engel et al., 2003) to account for deviations from multivariate normality and missing data. We assessed model fit through the (Satorra-Bentler scaled) chi-squared test, the comparative fit index (Robust CFI), the root mean square error of approximation (Robust RMSEA) with its confidence interval, and the standardized root mean squared residuals (Robust SRMR) (Schermelleh-Engel et al., 2003). Evaluation of model fit was defined as: Robust CFI (good fit > 0.97, acceptable fit 0.95-0.97), Robust RMSEA (good fit < 0.05, acceptable fit 0.05-0.08), and Robust SRMR (good fit < 0.05, acceptable fit 0.05–.10).

We followed the guidance set forth by McNeish and Wolf, 2020 for determining the proper weighting (equal or allowed to vary) and factorial structure (single or multiple factors) of our neighborhood crime data. To assess the optimal weighting of the neighborhood crime variables, we fit two types of factor models: parallel and congeneric. For the parallel factor model, we constrained all factor loadings (i.e., correlation between the observed crime indices and the Neighborhood Crime latent factor) to be equal. We also constrained the error (residual) variances for each neighborhood crime index to be equal. As a result, the parallel factor model assumes each neighborhood crime index contributes equally to the latent Neighborhood Crime construct. The second type of model we fit is known as a congeneric factor model. The congeneric factor model freely estimates the factor loading of each neighborhood crime index so that they are allowed to differ from each other. Thus, in contrast to the parallel factor model, the congeneric factor model does not assume that each neighborhood crime index contributes equally to the latent construct and, therefore, has a unique error variance (Graham, 2006). Furthermore, in this model, we constrained the variance of the latent variable, Neighborhood Crime, to 1 (McNeish and Wolf, 2020).

To compare the single-factor parallel and congeneric models, we used a likelihood ratio test, computing a scaled chi-squared difference test. We also compared the models’ Akaike information criterion (AIC, Bozdogan, 1987) and Bayesian information criterion (BIC, Bollen et al., 2014; Schwarz, 1978) estimates, which indicate whether increased model fit warrants increased model complexity (e.g., fewer degrees of freedom). We considered a model to be ‘better’ if the chi-squared difference test was significant (p<0.05), which indicated that the fits differed between models, and if the model’s AIC and BIC were lower than the model it was compared to. Depending on which single-factor model was determined to be better (parallel or congeneric), we repeated the SEM analyses with a corresponding two-factor model with the latent constructs Non-Violent Crime and Violent Crime. Here, the likelihood ratio test would assess whether the two-factor parallel or congeneric model fit better given its additional complexity (i.e., a second latent variable) compared to its single-factor counterpart. If none of the models displayed good or acceptable fit well to the data, we would not create a total crime sum score and would instead include each neighborhood crime index variable as a separate node in our bilayer network estimations. Note, the goal of this SEM analysis procedure was not to find the best-fitting model overall. Therefore, we did not fit any alternative models with more than two latent factors. Instead the aim was to establish whether unit-weighting (i.e., each crime variable contributes equally to the total crime sum score) and/or a single factor adequately represents the neighborhood crime data in this sample. This would indicate whether estimating the relationships of the individual crime indices is more appropriate than creating a new total crime sum score or multiple sum scores based on these six variables.

First, we describe the sum score analysis findings for the exposome-structural MRI data, which contained the same neighborhood crime index data (n = 170, 152 with crime data). The single-factor parallel model displayed poor fit to the data (*χ*^2^(19) = 296.348, p < .001, Yuan-Bentler scaling factor = .986, CFI = .531, RMSEA = .308 [.277 .339], SRMR = .148). Similarly, the single-factor congeneric model also displayed poor fit (*χ*^2^(9) = 132.484, p < .001, Yuan-Bentler scaling factor = 1.115, CFI = .791, RMSEA = .319 [.275 .364], SRMR = .088), although better than the parallel model (*χ*^2^Δ = 166.08, dfΔ = 10, AICΔ = 124.6, BICΔ = 94.4, p < .001). This suggests that, if sum scored, the neighborhood crime indices should not be unit-weighted (i.e., each crime variable contributes equally to the total crime sum score) for the exposome-brain structural MRI data. Furthermore, given the poor fit of the single-factor congeneric model, we did not create an aggregate sum score based on the Neighborhood Crime latent variable.

Next, we investigated the factorial structure of the exposome-structural MRI neighborhood crime index data to determine whether a two-factor model (Non-Violent Crime and Violent Crime) would show acceptable or good fit to the data. We fitted a congeneric two-factor model corresponding to the best fitting single-factor model. In line with the single-factor models, the two-factor congeneric model also fitted poorly to the crime data (*χ*^2^(8) = 120.107, p < .001, Yuan-Bentler scaling factor = 1.128, CFI = .813, RMSEA = .324 [.277 .373], SRMR = .093). However, it did outperform the single-factor congeneric model (*χ*^2^Δ = 12.100, dfΔ = 1, AICΔ = 10.2, BICΔ = 7.2, p < .001). Overall, these results indicate that the neighborhood crime indices from our exposome-structural MRI data should neither be unit-weighted nor combined into a single sum score. Therefore, we chose to include the individual neighborhood crime indices (aggravated assault, burglary, rape, larceny, murder, and robbery) as separate nodes to estimate their relationships in the exposome-cortical surface area and thickness networks.

We repeated the above sum score analyses for the exposome-functional MRI data, which contained a different set of neighborhood crime index data from the structural MRI sample (n = 130, of which 112 had crime index data). The single-factor parallel model exhibited poor fit to the data (*χ*^2^(19) = 239.228, p < .001, Yuan-Bentler scaling factor = .965, CFI = .557, RMSEA = .316 [.281 .352], SRMR = .158). Similarly, the single-factor congeneric model also fitted poorly to the data (*χ*^2^(9) = 110.357, p < .001, Yuan-Bentler scaling factor = 1.098, CFI = .767, RMSEA = .333 [.281 .388], SRMR = .096), although better than the parallel model (*χ*^2^Δ = 129.75, dfΔ = 10, AICΔ = 90.6, BICΔ = 62.6, p < .001). This suggests that, if sum scored, the neighborhood crime indices should not be unit-weighted (i.e., each crime variable contributes equally to the total crime sum score) for the exposome-functional MRI data. Moreover, since the single-factor congeneric model fitted poorly to the neighborhood crime data, we did not create an aggregate sum score based on the Neighborhood Crime latent variable.

Lastly, we replicated the previous investigation of the factorial structure of the neighborhood crime index data in exposome-functional MRI data. We fitted a congeneric two-factor model corresponding to the best fitting single-factor model. In line with the single-factor models, the two-factor congeneric model also fitted poorly to the crime data (*χ*^2^(8) = 96.202, p < .001, Yuan-Bentler scaling factor = 1.105, CFI = .796, RMSEA = .330 [.274 .390], SRMR = .099). However, it did outperform the single-factor congeneric model (*χ*^2^Δ = 14.282, dfΔ = 1, AICΔ = 12.8, BICΔ = 10.1, p < .001). These results suggest that, as with exposome-structural data, neighborhood crime index data for exposome-functional MRI data should neither be unit-weighted nor combined into a single sum score. Hence, we chose to use the individual indices for the exposome-participation coefficient and exposome-clustering coefficient multilayer network estimation.

